# Phase-separated stress granules and processing bodies are compromised in Myotonic Dystrophy Type 1

**DOI:** 10.1101/2021.06.14.448303

**Authors:** Selma Gulyurtlu, Monika S Magon, Patrick Guest, Panagiotis P Papavasiliou, Alan R Prescott, Judith E Sleeman

## Abstract

RNA regulation in mammalian cells requires complex physical compartmentalisation using structures thought to be formed by liquid-liquid phase separation. Disruption of these structures is implicated in numerous degenerative diseases. Myotonic Dystrophy Type 1 (DM1) is a multi-systemic trinucleotide repeat disorder resulting from a CTG expansion in the dystonia myotonica protein kinase gene (DMPK). The cellular hall-mark of DM1 is the formation of nuclear foci containing expanded DMPK RNA (CUGexp). We report here the deregulation of stress granules and processing bodies (P-bodies), two cytoplasmic structures key for mRNA regulation, in cell culture models of DM1. Alterations to the rates of formation and dispersal of stress granules suggest an altered ability to respond to stress associated with DM1, while changes to the structure and dynamics of stress granules and P-bodies suggest that a more widespread alteration to the biophysical properties of cellular structures may be a consequence of the presence of CUGexp RNA.

## Introduction

Myotonic Dystrophy Type 1 (DM1) is an autosomal dominant, multi-systemic disease (reviewed in Pettersson et al. (2015)) with an estimated global prevalence of 1:20000 (Theadom et al. (2014). Its symptoms are variable and can include myotonia, muscular atrophy, cardiac defects, cataracts and insulin resistance (Pettersson et al., 2015). The condition is caused by a CTG repeat expansion in the gene encoding Dystrophia Myotonica Protein Kinase (DMPK) on chromosome 19. Transcription of the mutant DMPK gene results in mutant transcripts containing a (CUG)_n_ repeat expansion in their 3’ untranslated region (3’UTR), which form an elongated metastable stem-loop structure (Pettersson & Aagaard, 2015). These transcripts accumulate to form nuclear foci (CUGexp foci) and recruit RNA-binding proteins (RBPs), notably Muscleblind-like Protein 1 (MBNL1), which interacts dynamically with the CUGexp foci showing rapid exchange with nucleoplasmic MBNL1 (Coleman et al., 2014; Ho et al., 2005). The similarities between DM1 and DM2, which results from a similar repeat expansion in an unrelated gene (Liquori et al., 2001), suggest that the nuclear CUGexp foci are the source of the pathology of DM1. The mechanisms by which the CUGexp foci result in cellular damage and the broad range of symptoms of DM1 are not understood, but are proposed to include disturbance of the normal cellular functions of MBNL1.

MBNL1 is the most abundant RNA metabolism regulator from the Muscleblind-like protein family, and is expressed in most tissues (Konieczny et al., 2014). MBNL1 has a key role as an alternative splicing regulator, known to act antagonistically with a second splicing regulator, CUG binding protein 1 (CUGBP1, also known as CELF1) (Junghans, 2009; Mahadevan et al., 2006). The splicing activity of CUGBP1 is increased in DM1, thought to be linked to the accumulation of MBNL1 in the CUGexp foci. Splicing alterations attributed to disturbance of the balance between MBNL1 and CUGBP1 activity have been documented in DM1. For example, MBNL1 inhibits Exon-5 inclusion in Cardiac Troponin T mRNA whereas CUGBP-1 promotes it (Wang et al., 2007). In contrast, Exon-11 inclusion in the Insulin Receptor (IR) is promoted by MBNL1, and inhibited by CUGBP1 (Savkur et al., 2001). However, both MBNL1 and CUGBP1 have diverse functions in regulating gene expression, in addition to their roles as alternative splicing regulators. MBNL1 is also implicated in RNA transport, stability and miRNA processing (Konieczny et al., 2014; Osborne et al., 2009; Rau et al., 2011; Wang et al., 2012) while CUGBP1 is implicated in mRNA translation, stability and decay (Masuda et al., 2012; Teplova et al., 2010).

Cellular RNA metabolism is complex, and its physical organisation within the cell involves the formation of RNA-rich bodies in the nucleus and the cytoplasm. In the nucleus, these include Cajal Bodies, Speckles and Paraspeckles, while in the cytoplasm, Processing Bodies (P-bodies) and stress granules play roles in RNA stability. These are all non-membranous and highly dynamic structures and there is evidence that at least some of them are formed by Liquid-Liquid Phase Separation (LLPS) (Banani et al., 2016; Jain et al., 2016; Kroschwald et al., 2015; Protter and Parker, 2016; Stanek and Fox, 2017). LLPS involves a high concentration of a molecule, or a mixture of molecules, forming a network of weak multivalent interactions resulting in a separate phase. This allows the structures to exhibit liquid-like properties, such as having a spherical, droplet-like morphology resulting from reduced surface tension and demonstrating fusion, fission and reversible disruption (Courchaine et al., 2016; Sawyer et al., 2019). Phase separated structures normally allow rapid exchange of components, which distinguishes them from the more solid protein aggregates (Markmiller et al., 2018; Stanek and Fox, 2017) that are implicated in pathologies including Amyotrophic Lateral Sclerosis (ALS) and Frontal Lobe Dementia (FTD) (Jain et al., 2016; Li et al., 2013; Markmiller et al., 2018; Ramaswami et al., 2013).

In mammalian cells, P-bodies and stress granules (Jain et al., 2016; Protter and Parker, 2016) are cytoplasmic structures, proposed to form by phase separation dependant on mRNA-protein interactions, with key roles in cellular signalling, metabolic machinery and the stress response (Anderson et al., 2015; Protter and Parker, 2016). They interact with each other both physically (Anderson et al., 2015; Protter and Parker, 2016; Shah et al., 2016), and functionally and also share a number of protein components, though the precise relationship between the two structures is not clear. P-bodies contain the RNA decay machinery, including decapping enzymes (DCP1/DCP2) and decapping activators (GE-1, EDC3, Par1, LSm1-7, RCK) (Anderson et al., 2015). P-bodies appear to be built on a scaffold of mRNA (Banani et al., 2016; Ditlev et al., 2018), with multivalent interactions between protein scaffolds and RNA being essential for P-body assembly (Ditlev et al., 2018). They have been specifically implicated in mRNA degradation, surveillance, translation repression and RNA-mediated gene silencing (Anderson et al., 2015; Protter and Parker, 2016). Stress granules form from mRNAs stalled in translation initiation, typically as a result of different forms of cellular stress such as oxidative stress, heat shock, hypoxia, starvation and viral infection (Anderson et al., 2015; Protter and Parker, 2016; Roy and Rajyaguru, 2018). Mammalian stress granules, in contrast to the smaller P-bodies, appear to contain a combination of liquid and more solid phases where different areas within the stress granule exhibit different levels of dynamic exchange of their components (Jain et al., 2016; Protter and Parker, 2016). In yeast, stress granules have a more solid structure (Ditlev et al., 2018; Jain et al., 2016; Kroschwald et al., 2015). On removal of stress, clearance of stress granules is mediated largely by a form of autophagy (Hardy et al., 2017; Mateju et al., 2017; Protter and Parker, 2016; Turakhiya et al., 2018) and the mRNPs sequestered within them during the stress response return to active translation (Lee and Seydoux, 2019). Aberrant persistent stress granules have been implicated in age-related (Mateju et al., 2017) and neurodegenerative diseases (Protter and Parker, 2016), such as ALS and FTD (Jain et al., 2016; Markmiller et al., 2018; Zhang et al., 2018). These are proposed to result from increased liquid-to-solid phase transitions within the stress granules (Mateju et al., 2017).

Increased cellular stress, resulting from the presence of CUGexp RNA foci has been suggested as a pathological effector in DM1 (Huichalaf et al., 2010; Ihara et al., 1995; Kumar et al., 2014; Toscano et al., 2005; Usuki and Ishiura, 1998; Usuki et al., 2000). Both MBNL1 and CUGBP1, key proteins implicated in the pathology of DM1, have been identified in stress granules in mammalian cells (Fujimura et al., 2008; Onishi et al., 2008), and increased incidence of stress granules reported in DM1 myoblasts compared to controls under normal growth conditions (Huichalaf et al., 2010). Furthermore, a defect in formation of stress granules in response to sodium arsenite has been reported in DM1 patient-derived fibroblasts expressing exogenous MyoD to mimic a myoblast phenotype, suggesting an impaired ability of DM1 cells to respond to stress (Ravel-Chapuis et al., 2016). We now identify both MBNL1 and CUGBP1 as components of P-bodies as well as stress granules in human eye lens epithelial cells and in a novel inducible cell model of DM1. Analysis of the dynamic behaviour of MBNL1 and CUGBP1 in live cells further implicates MBNL1 as a regulator of the turnover of cytoplasmic stress granules, suggesting a mechanism by which disruption of the sub-cellular distribution of MBNL1 by CUGexp RNA could result in deregulation of stress granules and, by extension, the fine control of cytoplasmic RNA metabolism.

## Material and Methods

### Plasmids, Transfection and Lentiviral Transductions

Plasmids pBI-Tet-CTG12 and pBI-Tet-CTG960 (Lee et al., 2012; Philips et al., 1998) were a gift from Prof. T Cooper, Baylor College of Medicine, Houston. These constructs contain a bidirectional tetracycline responsive promoter allowing the simultaneous expression of GFP and a DMPK1 mini-gene containing an interrupted CTG repeat length corresponding to a healthy control state (pBI-Tet-CTG12) or a disease-relevant expansion (pBI-Tet-CTG960). To generate plasmids pBI-Tet-CTG12_GFPMBNL1 and pBI-Tet-CTG960_GFPMBNL1, sequence and ligation independent cloning (SLIC) was used to introduce the GFP-MBNL1 cDNA sequence into the pBI-Tet-CTG12 vector after which the disease-relevant expansion was introduced by restriction digest and ligation. To generate plasmid pcDNA3rtTA3, the cDNA sequence encoding the third generation rtTA protein (rtTA3) was obtained by PCR from plasmid MXS_PGK::rtTA3-bGHpA (Addgene plasmid #62446) (Sladitschek and Neveu, 2015) using primers gctagtaagcttatgtctagactggacaagagcaaagtc and atcatggatccttacccggggagcatgtc and cloned into the pcDNA3 vector for mammalian expression.

The cDNA sequence for Dcp1a was sub-cloned from plasmid pEGFP-DCP1a-C1 (Aizer and Shav-Tal, 2008) a gift from Dr Y. Shav-Tal, Bar-Ilan University, Israel) into pmCherry-C1 using XhoI and EcoRI to form pmCherry-DCP1a-C1. Transient transfections were performed using Effectene Transfection reagent (QIAGEN), according to manufacturer’s instructions.

For lentiviral knockdown, oligonucleotides were based on Ambion siRNA sequences available from the Sigma MISSION shRNA library. For MBNL1 shRNA targeting the 3’UTR, sense sequence was CCGGGAGTAAAGGACGAGGTCATTACTCGAGTAATGACCTCGTCCTTTACTCTTTTTTG antisense sequence AATTCAAAAAAGAGTAAAGGACGAGGTCATTACTCGAGTAATGACCTCGTCCTTTACTC.

For CUGBP1 shRNA targeting the 3’UTR, sense sequence was CCGGCGTCAAGTACATCGTCCAAATCTCGAGATTTGGACGATGTACTTGACGTTTTTG antisense sequence AATTCAAAAACGTCAAGTACATCGTCCAAATCTCGAGATTTGGACGATGTACTTGACG. Oligonucleotide sequences were phosphorylated, annealed and then ligated into pLKO-Tet-on (Addgene plasmid #21915) (Wiederschain et al., 2009), to generate plasmids pLKO_MBNL1 and pLKO_CUGBP1. Lentiviral particles were produced using 293T cells and transduction of HeLa cells performed as described in (Wiederschain et al., 2009). Luciferase (control) plasmid was Tet-pLKO-puro_shLUC (Harwadt et al, 2016 (Harwardt et al., 2016), a gift from Dr Christina Paulus, University of St Andrews).

### Cell Culture

Human lens epithelial cells from DM1 patients (DMCat1-4) and age-matched controls (CCat1 and CCat2) have been described previously (Rhodes et al., 2006). These cells were grown at 37°C and 5% CO_2_ in Dulbecco’s Modified Eagle Medium (DMEM, Sigma) supplemented with 10% Foetal Bovine Serum (FBS, Sigma), 1% Penicillin/Streptomycin (Sigma) and 1% L-Glutamine (Sigma). HeLa cells were grown at 37°C and 5% CO_2_ in Dulbecco’s Modified Eagle Medium (DMEM, Sigma) supplemented with 10% Foetal Bovine Serum (FBS, Sigma), 1% Penicillin/Streptomycin (Sigma). Following transduction with lentiviral vectors, medium was supplemented with 250ng/ml Puromycin and 200μg/ml G418/Geneticin (Sigma).

Cell lines HeLa_CTG12_GFPMBNL1 and HeLa_CTG960_GFPMBNL1 were generated in two stages. HeLa cells were first transfected with the pcDNA3-rtTA3 construct and selected with G418. Individual clones were generated and screened for low background expression and high levels of induction of rtTA3. The best performing clone was co-transfected with either pBI-Tet-CTG12_GFPMBNL1 or pBI-Tet-CTG960_GFPMBNL1 and a linear puromycin marker (Invitrogen), selected with puromycin and clonal cell lines screened for inducible nuclear foci formation following addition of 1μg/ml doxycycline using fluorescence *in situ* hybridisation (Cy3-CAG10 probe) and fluorescence microscopy (GFPMBNL1). These cell lines were maintained as for the parental HeLa cell line, with G418 and puromycin added periodically to ensure retention of the added sequences. When required, expression of the constructs was induced using 1μg/ml doxycycline for 24, 48 or 72 hrs as indicated.

### Western Blotting

Cells were lysed with ice-cold nuclear lysis buffer (50mM Tris-HCl pH7.5; 0.5M NaCl; 1% Igepal; 1% Sodium deoxycholate; 0.1% SDS; 2mM EDTA; cOmplete Protease Inhibitor tablet (Roche)) for 5 minutes, then passed through a QIAshredder (QIAGEN) for 2 minutes at 13000RPM centrifugation. Lysates were electrophoresed in a 10% Acrylamide gel and transferred onto a nitrocellulose membrane (GE Healthcare) using a semi-dry blotter (BioRad). 5% skimmed milk powder in PBS was used as blocking solution before detection with primary antibodies Mbla (mouse anti-MBNL1 (Holt et al., 2007) 1:50); rabbit anti-GFP (Abcam ab290 1:500); rabbit anti-CUGBP1 (Abcam ab129115 1:1000) and secondary antibodies goat anti-rabbit-AlexaFluor790 and goat-anti-mouse-AlexaFluor680 (Invitrogen 1:20000). Blots were imaged using a Licor Odyssey CLx.

### Immunocytochemistry

Cells were seeded on 18mm square glass coverslips and incubated in normal DMEM for 24 hours minimum. If required, 0.5μM NaAsO2 was added for 45 minutes before fixation followed by 5% 1,6-Hexanediol in water for 20 minutes before fixation, where appropriate. Fixation was performed using 3.7% Paraformaldehyde (PFA) (Sigma) in PHEM buffer (60mM PIPES, 25mM HEPES, 10mM EGTA, 2mM MgCl2, pH6.9) for 10 minutes at room temperature. Immunostaining was carried out essentially as described in (Sleeman et al., 2003), with 1% goat serum in PBS as a blocking buffer and 1 hour incubations with antibodies. Primary antibodies were Mbla mouse anti-MBNL1(Holt et al., 2007) 1:15; rabbit anti-TIA1 (Proteintech 12133-2-AP) 1:50; rabbit anti-CUGBP1 (Abcam ab129115, 1:100); rabbit anti-GE1 (Cell Signalling Technology #2548, 1:400). Secondary antibodies were goat-anti-rabbit AlexaFluor594, goat-anti-mouse AlexaFluor488, goat anti-mouse and goat-anti-rabbit Cy5 (All Jackson Laboratories, 1:250). 1.6μg/ml DAPI (Sigma Aldrich) in water was used to stain the nucleus. Coverslips were mounted using ProLong Gold Antifade (Invitrogen).

### RNA Fluorescence In Situ Hybridisation (FISH)

Cells were seeded on 18mm square glass coverslips and incubated in normal DMEM or DMEM supplemented with 1μg/ml Doxycycline as appropriate. Cells were fixed using 3.7% Paraformaldehyde (PFA) (Sigma), in DEPC-treated PBS and permeabilised with 70% ethanol in DEPC-treated water for 1 hour. A Cy3 labelled CAG10 DNA probe (5’-/5Cy3/(CAG)_10_, integrated DNA technologies) was diluted in hybridisation buffer (2X SSC; 10% w/v dextran sulfate; 10% formamide) to a final concentration of 200nM and incubated with the samples for 4 hours at 37°C. Samples were washed with 2X SSC, 3 × 5 minutes. 1.6μg/ml DAPI (Sigma Aldrich) in water was used to stain the nucleus. Coverslips were mounted using ProLong Gold Antifade (Invitrogen).

### Microscopy and Image Processing

Fixed samples were imaged using a 1.35na 100X objective on an Olympus DeltaVision RT microscope (Applied Precision) with 2×2 binning used for weaker samples and 0.2μm sectioning for *z*-stacks. Exposure times using DAPI, FITC, TRITC and Cy5 filter sets were chosen to aim for maximum intensities of 3600. Deconvolution, where indicated, was carried out using Volocity 6.3 image analysis software (Quorum Technologies) with calculated point-spread functions.

### High Resolution Image capture and Analysis

Images were captured using the Zeiss LSM 880 laser scanning microscope using the Airyscan detector with either the x63 or x100 objectives. This detector increases the resolution of the final image from a theoretical maximum of 0.2 μm to 0.1 μm (Huff, 2015). Correlation parameters were collected from the images using a custom programme written in the Volocity quantitation module (Quorum Technologies): GFP-MBNL1 positive structures were identified using the Otsu method (Otsu, 1979), objects in the nucleus were removed by identifying those intersecting with DAPI labelled regions. All remaining cytoplasmic structures were interrogated for the correlation between GFP-MBNL1 and CUGBP1 intensities.

### Live Cell Imaging

Cells were seeded onto 40mm round coverslips (Intracel) for 24 hours before transfections or transductions (where appropriate) were performed. After an additional 24 hours, the coverslips were placed into a POC-R imaging chamber (Zeiss) within an environmental incubator (Solent Scientific) on an Olympus DeltaVision RT microscope (Applied Precision), with a quantifiable laser module including 488 nm and 405 nm lasers, and either maintained at 37°C with 5% CO_2_ or incubated in CO_2_-independent medium (Life Technologies). *Z*-stacks of images separated by 500 nm were collected at a minimum of 3-min intervals using a 1.35 na 100X objective. Exposure times were optimized to give a maximum intensity value of ~1000 in each *z*-stack or in the first timepoint of each time-lapse series.

### Fluorescence Recovery After Photobleaching (FRAP)

FRAP was carried out using an Olympus DeltaVision RT microscope using a 1.35NA 100X objective, with the chamber heated to 37°C. A 488nm laser was used to photo-bleach GFP, at 100% power (2.5s). A time-course using the FITC filter was taken with 3 images before the laser fire and 25 images post-fire, using adaptive time intervals as implemented in the DeltaVision software. Where appropriate, P-bodies were identified using pmCherry-Dcp1A, viewed through the TRITC filter before FRAP. Image analysis was carried out using Volocity 6.3 image analysis software (Quorum Technologies). The photobleached region was indicated by a point and the intensity value measured against time. The values were corrected for background signal and photobleaching, and normalised by setting the intensity immediately after the firing event as 0%, and the highest of the pre-fire images as 100%. A one-phase association curve analysis was used in Prism 8 (GraphPad) to determine the mobile fraction and half-time of recovery for each data set.

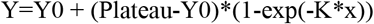

### Statistical Analyses

All statistical analyses, including ANOVA, T-tests, Mann-Whitney, D’Agostino-Pearson omnibus K2 normality test and removal of outliers as appropriate were carried out using Prism 8 (GraphPad). For analyses where all relevant datasets showed normal distribution, a one-way ANOVA test was used. For analyses where some, or all, datasets failed a D’Agostino-Pearson omnibus K2 normality test, a Mann-Whitney test was used.

## Results

The hallmark cellular defect in DM1 is the presence of CUGexp RNA foci formed by mutant DMPK transcripts in the cell nucleus. While some DMPK mRNA is clearly exported to the cytoplasm and translated (Gudde et al., 2017), the global effect of the presence of CTGexp RNA and formation of nuclear CUGexp foci on cellular RNA metabolism is not understood, although the sequestration of the multi-functional RBP, MBNL1, within the foci is thought to be a major cause of DM1 pathology. Lens cataract is the most prevalent symptom in DM1 (Romeo, 2012; Smith and Gutmann, 2016). We have previously determined that the CUGexp foci in human lens epithelial cells (HLECs) derived from DM1 patients contain only 0.2-0.5% of cellular MBNL1 (Coleman et al., 2014), making sequestration of MBNL1 within the nucleus an unlikely explanation for DM1-associated cataract. The transparency of the lens is achieved by the differentiation of a stem cell pool within the lens epithelium, during which transcription is shut down and all cellular organelles and inclusions are lost (Dahm and Prescott, 2002; Gribbon et al., 2002). Lens formation relies on forms of autophagy: the same process that is responsible for clearance of stress granules under certain conditions (Hardy et al., 2017; Mateju et al., 2017; Protter and Parker, 2016; Turakhiya et al., 2018). Together with evidence for alteration of stress granules in DM1 and our previous observation that MBNL1 localises to stress granules following transcriptional arrest in HLECs (Coleman et al., 2014), this led us to investigate the presence of the DM1-associated proteins MBNL1 and CUGBP1 in cytoplasmic structures involved in RNA metabolism.

### Muscleblind-Like Protein 1 (MBNL1) and CUG-Binding Protein 1 (CUGBP1) colocalise in P-bodies in human lens epithelial cells

Our initial examination of the sub-cellular distributions of endogenous CUGBP1 and MBNL1 in HLECs revealed a striking contrast between their distributions in the nucleus, where MBNL1 localises to CUGexp foci and CUGBP1 does not, and the cytoplasm, where the two proteins co-localise in discrete punctate structures (Fig.1A). Further analysis of these structures demonstrated them to be P-bodies, as both MBNL1 and CUGBP1 co-localise with the PB marker GE1 (Fig.1A). While accumulation of CUGBP1 has been reported in P-bodies (Yu et al., 2013), MBNL1 has not previously been reported to accumulate in these structures. Our observation of both of these proteins in P-bodies points to close links between the cytoplasmic functions of MBNL1 and CUGBP1.

**Figure 1.**
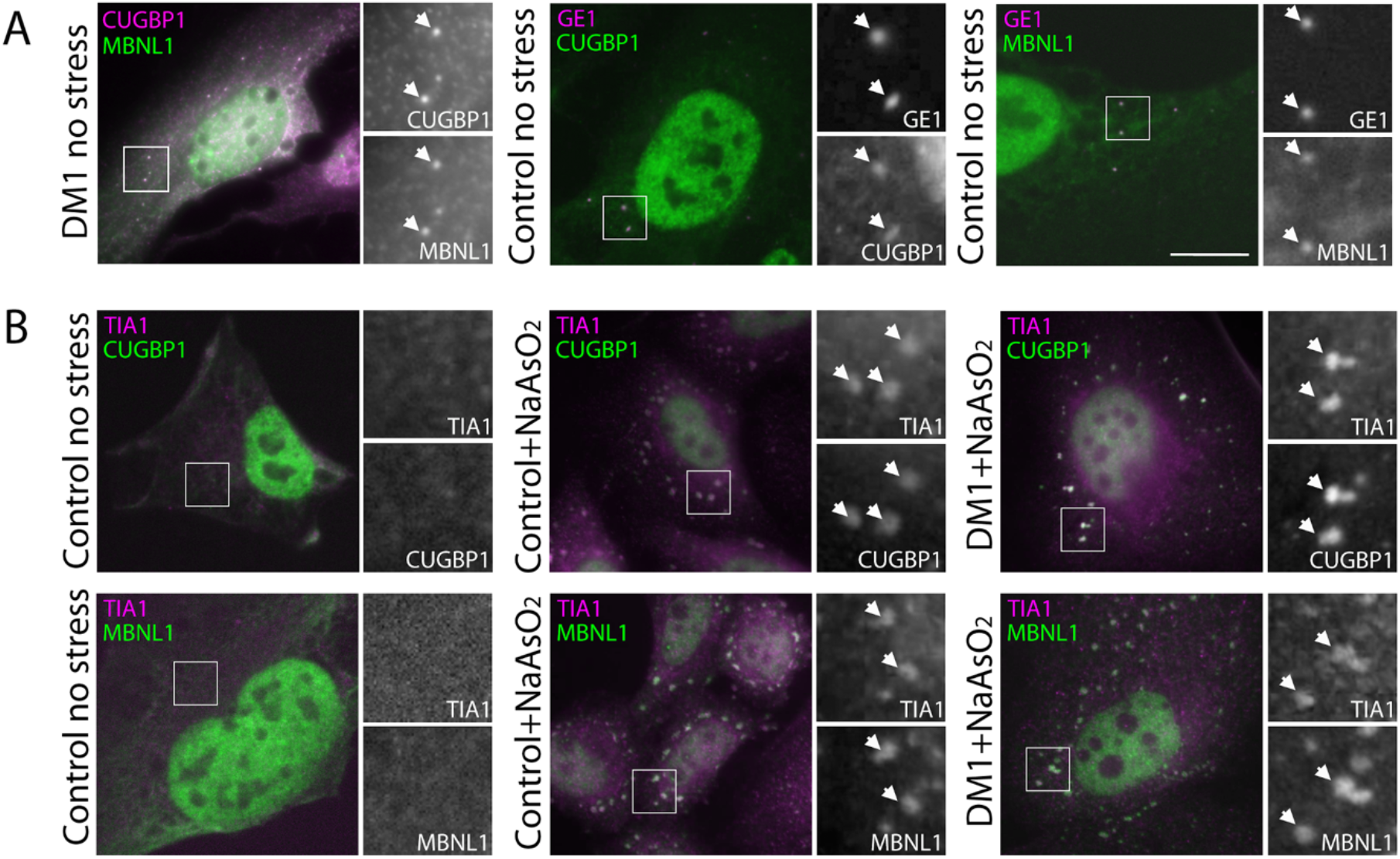
MBNL1 and CUGBP1 both localise to Processing Bodies (P-bodies) in Human Lens Epithelial cells and re-localise to stress granules following treatment with sodium arsenite. A) endogenous MBNL1 (left panel, green on overlay) and CUGBP1 (left panel, magenta on overlay) co-localise in punctate cytoplasmic structures (arrows) despite showing little co-localisation in the nucleus. Counter-staining with antibodies against the P-body marker, GE1 (centre and right panels, magenta on overlay), demonstrates that both CUGBP1 (centre panel, green on overlay) and MBNL1 (right panel, green on overlay) localise to P-bodies (arrows). This is seen in DM1 and control cell lines. B) Stress granules, detected with antibodies against TIA1 (magenta on overlay) are not seen in unstressed cells (left panels). Following treatment with 0.5mM sodium arsenite for 45 mins, control cells (centre panels) and DM1cells (right panels) show clear cytoplasmic stress granules (arrows) identified by antibodies against TIA1 (magenta on overlays) that contain both CUGBP1 (top row green on overlays) and MBNL1 (bottom row, green on overlays) Bar = 7μm.

### MBNL1 and CUGBP1 both re-localise to stress granules in human lens epithelial cells following treatment with sodium arsenite

MBNL1 and CUGBP1 have both been reported to accumulate in stress granules (Fujimura et al., 2008; Kedersha and Anderson, 2009) in different cell types using varied strategies to induce cellular stress. Furthermore, myoblasts from DM1 patients have been reported to display stress granules under normal growth conditions, suggesting a higher basal level of stress compared to controls (Huichalaf et al., 2010). While we saw no evidence of stress granules in the DM1 HLEC lines under normal growth conditions (Fig.1A), the close relationship between P-bodies, where both proteins co-localised, and stress granules led us to investigate the effects of stress on the cytoplasmic distribution of MBNL1 and CUGBP1. We used sodium arsenite (NaAsO2) to induce stress granule formation in HLECs. Sodium arsenite induces stress granules through oxidative stress by activating eIF2αK (Aulas et al., 2017). Using TIA1 as a marker for stress granules, we showed that stress granules form readily in HLECs in response to sodium arsenite and that MBNL1 and CUGBP1 both accumulate in stress granules (Fig.1B).

### Stress granules show no delay in formation but disperse more quickly in DM1 human lens epithelial cells compared to controls

Having shown that stress granules are not present in DM1 HLECs under normal growth conditions, suggesting no increase in basal stress levels, and that stress granules containing MBNL1 and CUGBP1 form readily in response to sodium arsenite, we next sought to investigate the kinetics of stress granule formation and loss in DM1 HLECs compared to controls to determine whether DM1 HLECs are more sensitive to cellular stress. Cells fixed after differing lengths of treatment with sodium arsenite showed no delay in formation of stress granules, detected using endogenous TIA1, in DM1 HLECs compared to control HLECs (Fig.2A). In contrast, cells pre-treated with sodium arsenite to induce stress granules then allowed to recover showed a more rapid loss of stress granules, detected using endogenous TIA1 and MBNL1, from DM1 cells compared to controls (Fig.2B). This suggests that DM1 cells differ from controls in their ability to recover from stress.

**Figure 2.**
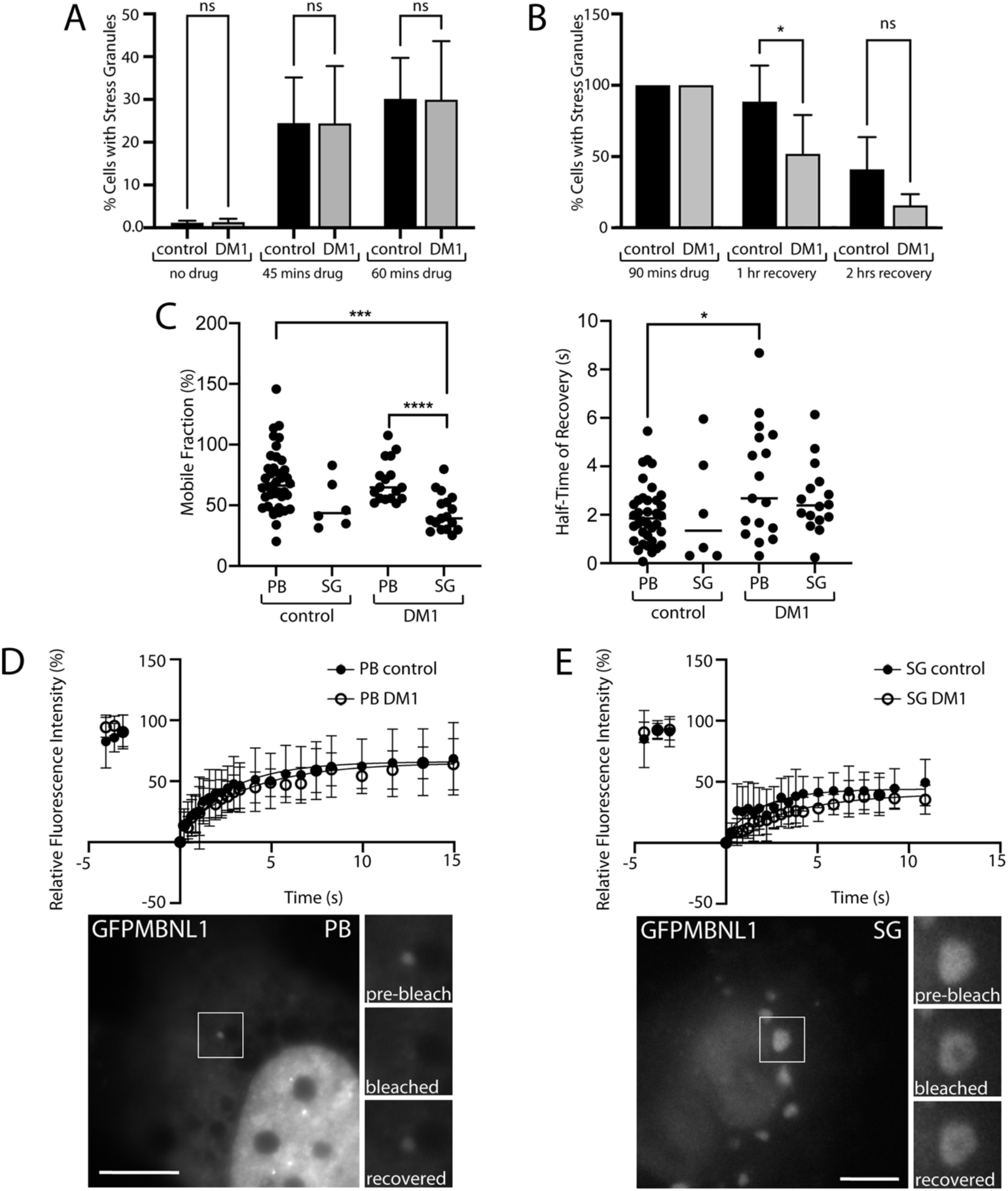
Kinetic analysis of Stress granules and P-bodies reveals an increased rate of loss of Stress Granules and slightly altered MBNL1 dynamics in DM1 HLECs compared to controls. A) Graph of percentage of cells containing stress granules detected with antibodies to endogenous TIA1 shows no significant difference in their formation between DM1 cells and controls following 45 or 60 minutes treatment with 0.25mM sodium arsenite. (n=3015 cells pooled from 3 cell lines for DM1 and 1776 cells pooled from 2 cells lines for controls from 2 independent experiments). B) Graph of percentage of cells containing stress granules detected with antibodies to endogenous TIA1 and MBNL1 following recovery from sodium arsenite treatment of 1hr or 2hrs, normalised to the percentage of cells showing stress granules following 90 minutes of drug treatment. Significantly fewer DM1 cells show stress granules remaining at the 1hr time point compared to controls. (n=1086 cells pooled from 3 cell lines for DM1 and 708 cells pooled from 2 cell lines for controls from 2 independent experiments. Data shown as mean +/-SD *P<0.05. C) Graphs showing the Mobile Fraction (left) and Half-Time of fluorescence Recovery (right) of GFPMBNL1 in P-bodies (PB) and stress granules (SG) in Control and DM1 HLECs.*P<0.05; ***P<0.001; ****P<0.0001. D and E) Curve fit of relative fluorescence recovery of MBNL1-GFP over time following bleaching using a 488nm laser at 2.5sec pulse in P-bodies (PB) (D) and stress granules (SG) (E), together with representative images from the experiment. The structure bleached in each case is highlighted by a white box with an expanded view shown before the bleach, immediately after the bleach and following recovery. Bar = 7μm. Closed circles on graphs indicate P-bodies/stress granules in control cells (CCat1 and CCat3, n=39 total for P-bodies and 6 for stress granules); open circles indicate P-bodies/stress granules in DM1 cells (DMCat1, DMCat2 and DMCat4, n=17 total for P-bodies and 17 for stress granules). All data displayed as mean +/- SD. See also figure S1.

### MBNL1 Dynamics are unaffected in stress granules in DM1 human lens epithelial cells and altered in P-bodies

We next used a fluorescence recovery after photobleaching (FRAP) approach to examine the dynamics of MBNL1 in stress granules and P-bodies, using transiently expressed GFP-MBNL1, previously validated to behave similarly to endogenous MBNL1 in these cell lines (Coleman et al., 2014). Previous studies have shown that in mammalian cells, these two related structures are both highly dynamic, displaying the properties of liquid droplets (Kroschwald et al., 2015). Persistent stress granules have long been associated with degenerative conditions, notably ALS (Li et al., 2013) but the dynamics of MBNL1 within stress granules and P-bodies in the context of DM1 has not been examined.

To analyse the dynamics of stress granules, GFPMBNL1 was transiently expressed in HLECs for 24hrs prior to incubation for 45 minutes with 0.5mM sodium arsenite. A diffraction-limited spot of laser excitation was then used to bleach GFPMBNL1 in stress granules and the subsequent recovery fitted to a one-phase exponential recovery model (Fig.2C and E). To analyse the dynamics of P-bodies, both GFPMBNL1 and mCherry-Dcp1a (a marker for P-bodies) were transiently expressed for 48hrs prior to FRAP experiments to enable unambiguous identification of P-bodies (Fig.2 C and D and supplementary figure S1). This modelling (Fig.2C) showed no statistically significant changes in the mobile fraction or halftime of recovery of GFPMBNL1 in stress granules in DM1 cells compared to controls suggesting that stress granules, once formed, are not defective in DM1 cells in terms of MBNL1 dynamics. The mobile fraction of GFPMBNL1 in P-bodies in both DM1 and control cells was significantly higher than that for stress granules in DM1 cells (69.6%+/-17.2 for PBs in DM1, 70.1%+/-25.2 for PBs in control, 43.9%+/-15.2 for SGs in DM1, 50.6% +/-20.2 for SGs in control), suggesting that GFPMBNL1 has a more dynamic relationship with P-bodies than with stress granules overall. Furthermore, a significant increase in the half-time of recovery for GFPMBNL1 in P-bodies was seen for DM1 cells compared to controls (3.4s+/2.3 for DM1, 2.0s+/-1.2 for control), which might indicate an altered relationship of MBNL1 with P-bodies associated with DM1.

### Simultaneous expression of a DMPK mini-gene with GFPMBNL1 models DM1 cells with CUGexp foci that can be detected in living cells

Patient-derived HLECs are a highly relevant cell model in which to study the cellular pathology of DM1. However, the use of patient cell lines comes with the key drawback that the lines are of diverse genetic origins making age, severity of disease and length of CUG repeats different between them. In order to be able to assess the effect of the CUGexp repeats on stress granule structure and dynamics directly, we generated an inducible HeLa cell model. These stable HeLa cell lines contain a bi-directional tetracycline responsive promoter driving simultaneous expression of a DMPK mini-gene and GFPMBNL1. To mimic the DM1 phenotype, the DMPK mini-gene contains an interrupted CTG repeat length of 960 (HeLa_CTG_960__GFPMBNL1 or CTG960). As the disease only manifests from >50 repeats (Lee et al., 2012) in the control cell line the DMPK mini-gene contains only 12 repeats (HeLa_CTG_12__GFPMBNL1 or CTG12).

Fluorescence microscopy and immunoblotting (Fig.3) confirmed that no detectable GFPMBNL1 was expressed in the absence of doxycycline (Dox, a tetracycline equivalent), while addition of Dox resulted in expression of GFPMBNL1 in both cell lines. In the control cell line, HeLa_CTG_12__ GFPMBNL1, the characteristic sub-cellular distribution of endogenous MBNL1 localised in the nucleus and, more weakly, in the cytoplasm was seen. In the DM1 model cell line, HeLa_CTG_960__GFPMBNL1, clear nuclear accumulations of GFP-MBNL1 were seen (Fig.3A). These were confirmed as CUGexp foci using RNA fluorescence *in situ* Hybridisation (FISH) with a Cy3-CAG_10_ probe against the trinucleotide repeats, allowing nuclear GFPMBNL1 accumulation to be used as a marker for CUGexp foci in subsequent fixed and live cell experiments. Comparison of the signals for endogenous and GFP-tagged MBNL1 detected with anti-MBNL1 by immunoblotting reveals a moderate over-expression of GFP-tagged MBNL1 in both lines.

**Figure 3.**
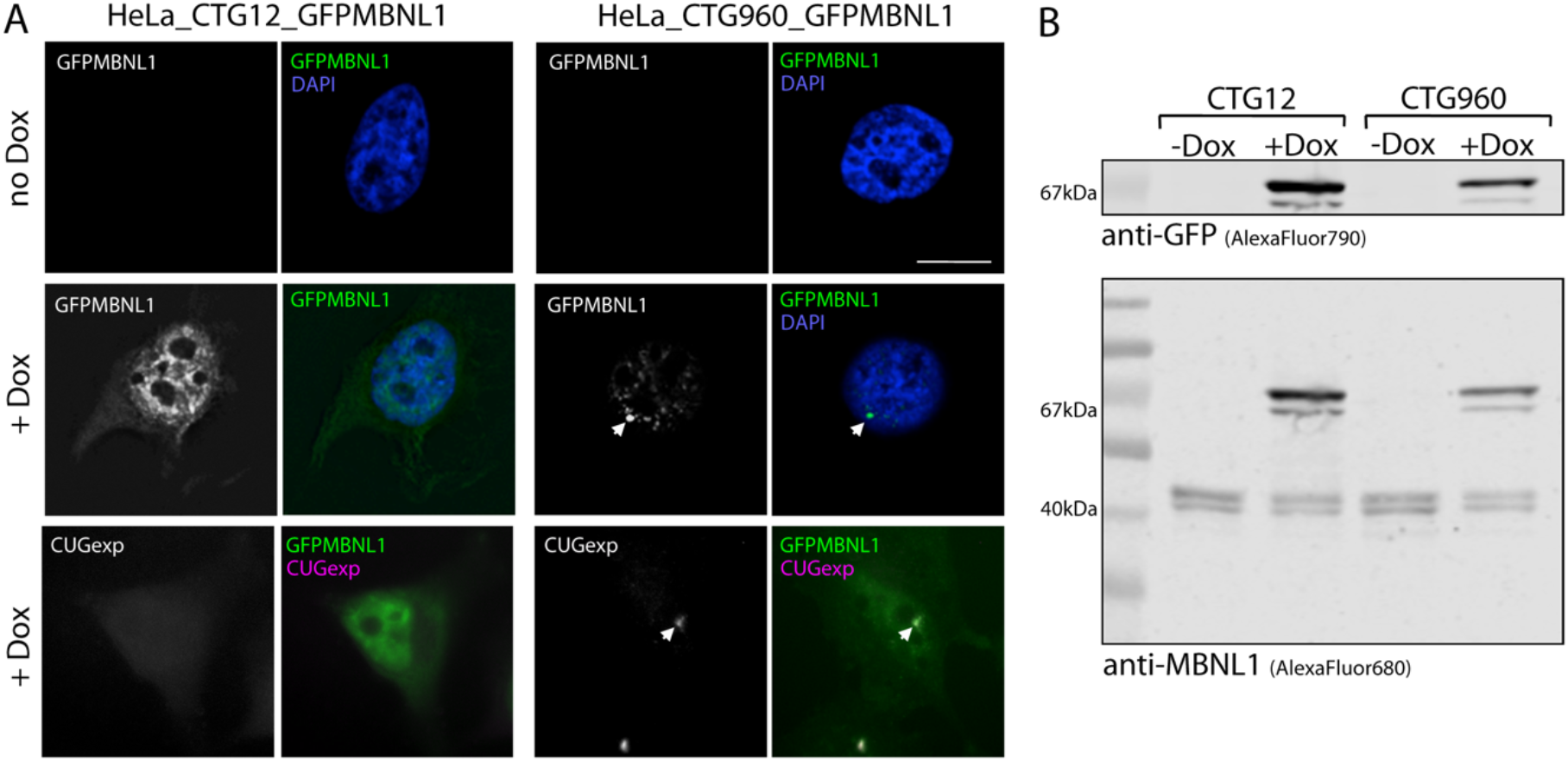
Simultaneous expression of a DMPK mini-gene with GFPMBNL1 models DM1. A) HeLa cells containing a DMPK minigene with 12 (left) or 960 (right) CTG trinucleotide repeats and GFPMBNL1 under a bi-directional tetracycline responsive promoter. Cells grown in the absence of Dox (top row) show no expression of GFPMBNL1 in the cytoplasm or nucleus (counterstained with DAPI, blue on overlay). Cells induced with 1μg/ml Doxycyline for 48hrs show clear expression of GFPMBNL1 (middle row, green on overlay), forming distinct nuclear foci (arrows) in the CTG960 line and the characteristic localisation of GFPMBNL1 to the nucleus and, more weakly, the cytoplasm in the CTG12 line. RNA FISH using a probe against the CUGexp RNA (bottom row, magenta on overlay) confirms that the GFPMBNL1 nuclear foci (arrows) are genuine CUGexp foci in the CTG960 line (right) and that no CUGexp RNA is detected in the CTG12 line (left). Bar = 10μm. B) Immunoblot detection of PAGE-separated whole-cell lysates using antibodies against GFP (top, AlexaFluor790) and MBNL1 (bottom, AlexaFluor680) confirm successful expression of GFPMBNL1 (67KDa) in both the CTG12 line (left) and the CTG960 line (right) in cells treated with Dox for 48hrs (+Dox) but not in uninduced cells (-Dox). All samples show the characteristic double-band just above the 40kDa marker for endogenous MBNL1 (bottom).

### MBNL1 and CUGBP1 co-localise in P-bodies and sub-regions of stress granules in HeLa models of DM1, with subtle changes in their co-localisation associated with the presence of CUGexp foci

In un-stressed cells from lines HeLa_CTG_12__GFPMBNL1 and HeLa_CTG_960__GFPMBNL1, P-bodies are less numerous and less distinct than in HLEC cells (Fig.4A). Those that are present accumulate small amounts of both GFPMBNL1 and CUGBP1 in the control HeLa_CTG_12__GFPMBNL1 line, but P-bodies in the DM1 model line, HeLa_CTG_960__GFPMBNL1 do not contain detectable amounts of GFPMBNL1. This suggests that P-body components are altered by the presence of the CTG960 expansion. Following treatment with sodium arsenite to induce cellular stress, both cell lines showed increased numbers of prominent P-bodies, detected with GE1, in addition to clear stress granules containing both GFPMBNL1 and CUGBP1 (Fig.4B). Using super-resolution airyscan microscopy, the localisation of both GFPMBNL1 and CUGBP1 was revealed to be non-uniform within each stress granule and largely co-incident (Fig.4C). In both cell lines the co-localisation between the two proteins, measured using Pearson’s co-efficient, was very high. However, a statistically significant increase seen in the HeLa_CTG_960__GFPMBNL1 cell line compared to the HeLa_CTG_12__GFPMBNL1 control line (Fig.4D) (0.90+/-0.05 for CTG960; 0.87+/-0.07 for CTG12), suggests a subtle change in stress granule architecture associated with the presence of CUGexp RNA. In arsenite-treated cells, some P-bodies in line HeLa_CTG_960__GFPMBNL1 do contain amounts of GFPMBNL1 detectable by airyscan microscopy, but the amount of GFPMBNL1 in P-bodies as a percentage of total cellular GFPMBNL1 is smaller in comparison with the control line, HeLa_CTG_12__GFPMBNL1 (Fig.4E) (0.08%+/-0.1 for CTG960; 0.1%+/-0.1 for CTG12) and supplementary figure S2.

**Figure 4.**
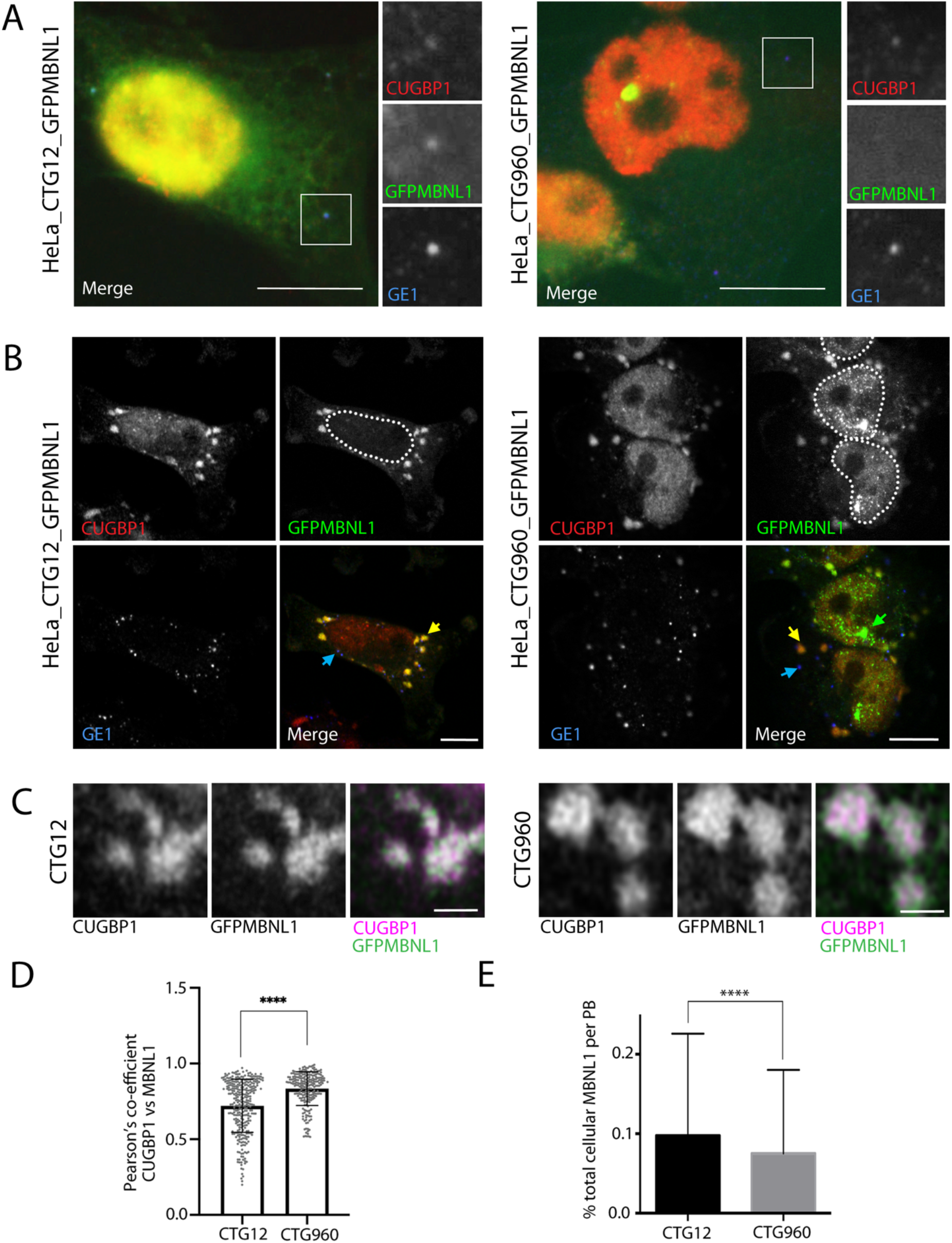
MBNL1 and CUGBP1 co-localise in P-bodies and sub-regions of stress granules in HeLa models of DM1. A) In cells from line HeLa_CTG_12__MBNL1-GFP (left) CUGBP1 (red on merge) and GFPMBNL1 (green on merge) co-localise with GE1 (blue on merge) in cytoplasmic P-bodies (expanded view is of PB highlighted by white box on overlay). In cells from line HeLa_CTG_960__MBNL1-GFP (right), CUGBP1 (red) colocalises with GE1 (blue) in P-bodies, but GFPMBNL1 (green) is not detectable (expanded view is of PB highlighted by white box on merge). Bar=10μm. Red and green signals in these images have been adjusted to visualise the cytoplasmic P-bodies, resulting in saturation of the nuclear signal. B) CUGBP1 (red on merge) and GFPMBNL1 (green on merge) co-localise in stress granules (yellow arrows) in cells from lines HeLa_CTG_12__GFPMBNL1 and HeLa_CTG_960__MBNL1-GFP following arsenite treatment. Large number of P-bodies (blue arrows), detected with GE1 (blue on merge) are also seen in both cell lines. Nuclear foci of GFPMBNL1 (green arrows) are clearly seen in the CTG960 line only. Dotted lines on GFPMBNL1 images show the approximate outlines of nuclei. Bar = 10μm. C) Super-resolution airyscan images of stress granules showing non-uniform distribution of CUGBP1 (magenta) and GFPMBNL1 (green). Bar=1μm. D) Graph of pearson’s co-efficient of co-localisation between CUGBP1 and GFPMBNL1 in stress granules (****P<0.00001, n=204 stress granules in CTG960 and 229 stress granules in CTG12 from 3 independent experiments). E) Graph of percentage of total cellular GFPMBNL1 found in P-bodies following sodium arsenite treatment in cells from lines HeLa_CTG_12__GFPMBNL1 and HeLa_CTG_960__GFPMBNL1 n=924 P-bodies in CTG12 cells and 670 P-bodies in CTG960 cells, ****P<0.0001. All data displayed as mean +/- SD. See also figure S2.

### Stress granule formation and dispersal are altered by CUGexp foci formation in an inducible HeLa cell model

To investigate further the effects of the presence of CUGexp foci on stress granules, we used time-lapse imaging of cells during response to and recovery from sodium arsenite treatment. Cell lines HeLa_CTG_960__GFPMBNL1 and HeLa_CTG_12__GFPMBNL1 were treated with doxycycline for 24hrs to induce expression of the DMPK mini genes (with and without extended CTG repeats) and GFPMBNL1. Treatment with sodium arsenite was then carried out under time-lapse microscopy, with Z-stacks of images taken every 4 minutes until stress granule formation was clearly seen (Fig.5A). This revealed a pronounced delay in formation of stress granules in cells containing CUGexp foci (HeLa_CTG_960__ GFPMBNL1, 36 min +/-12) compared to those without (HeLa_CTG_12__ GFPMBNL1, 15 min +/- 2) (Fig.5B). As cellular stress induced by sodium arsenite is reversible and dispersal of stress granules at least as important for their regulation as is their formation, we next investigated the timing of dispersal of stress granules in HeLa_CTG_960__MBNL1-GFP compared to controls. Cells from lines HeLa_CTG_960__ GFPMBNL1 and HeLa_CTG_12__ GFPMBNL1 were induced with doxcycline for 24 or 72 hours then treated for 45 minutes with 0.5mM sodium arsenite to ensure full induction of stress granules in both lines. The sodium arsenite was then thoroughly rinsed away and the cells imaged during their recovery. Cells were marked as recovered when no clear stress granules remained (Fig.5A). In contrast to the delay in formation of stress granules associated with CUGexp foci, stress granule dispersal occurred more quickly in cells containing CUGexp foci (HeLa_CTG_960__ GFPMBNL1, 133 mins +/-38 at 24 hrs and 79 mins +/- 11 at 72 hrs) compared to those lacking them (HeLa_CTG_12__ GFPMBNL1, 158 mins +/-30 at 24 hrs and 159 mins +/- 28 at 72 hrs). The increase in speed of dispersal was more pronounced after 72 hrs than after 24 hrs (Fig.5B). The disassembly seen using MBNL1 as a marker for stress granules closely resembles that reported previously, using GFP-G3BP1 as a marker (Wheeler et al., 2016), with breakdown of large stress granules into much smaller fragments (supp movie 2), suggesting that we are visualising the disassembly of stress granules, rather than simply the loss of MBNL1 from stress granules. Taken together, these data show that the presence of the CUG expansion affects both stress granule formation in response to sodium arsenite treatment and their dispersal on removal of the stress, suggesting an overall defect in stress granule regulation associated with the presence of the characteristic CUG expansion of DM1.

**Figure 5.**
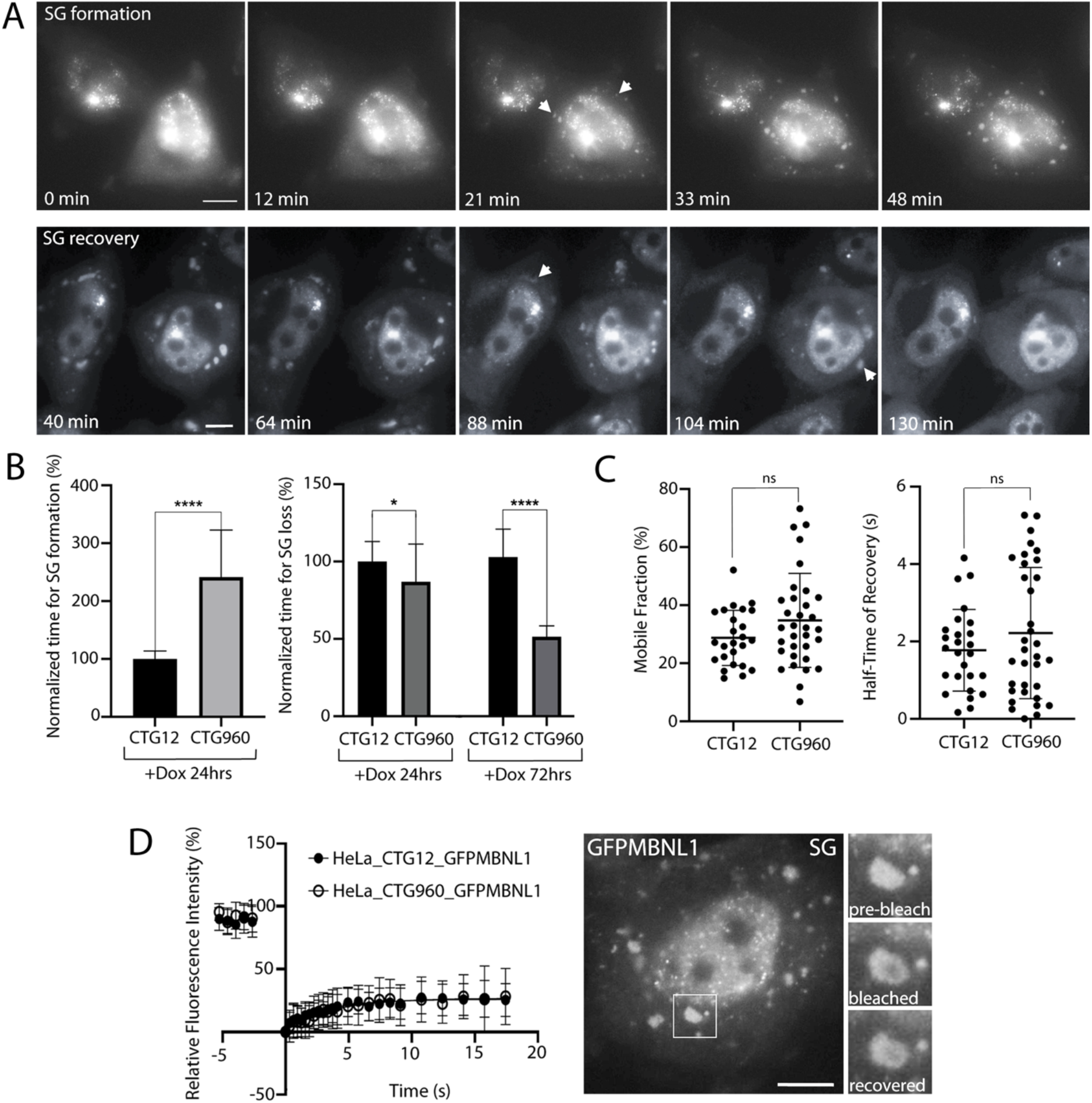
Kinetic analysis of stress granules reveals altered stress granule formation and dispersal in an inducible model of DM1 and increased variance in MBNL1 dynamics. A) Selected time-points from representative time-lapse sequences showing formation (top) and dispersal (bottom) of stress granules (SG, arrows) in the CTG960 cell line. The ‘formation time’ of stress granules was recorded as the time-point in which numerous stress granules could first be detected: 21 minutes for the example shown. The time of loss of stress granule formation was recorded as the first time-point in which clear stress granules could no longer be detected (108 minutes for the example shown). Bar=10um. See also movies S1 to S3. B) Graph of time taken for stress granule formation after 24 hrs of Dox induction (left) and SG loss after 24 hrs and 72 hrs of Dox induction (right), normalised to the mean values seen in control (CTG12) cells (n=19 CTG12 and 27 CTG960 cells for stress granule formation P<0.0001; n=35 CTG12 and 25 CTG960 cells for stress granule loss at 24 hrs P<0.05; n=33 CTG12 and 18 CTG960 cells for stress granule loss at 72 hrs P<0.0001, each experiment is representative of 2 independent experiments C) Graphs showing the Mobile Fraction (left) and Half-Time of fluorescence Recovery (right) of GFPMBNL1 in stress granules in CTG960 and CTG12 HeLa cells. D) Curve fit of relative fluorescence recovery of GFPMBNL1 over time following bleaching using a 488nm laser at 2.5sec pulse in stress granules, together with representative images from the experiment. The stress granule bleached is highlighted by a white box with an expanded view shown before the bleach, immediately after the bleach and following recovery. Bar = 7μm. Closed circles indicate stress granules in HeLa_CTG_12__GFPMBNL1 cells (n=41); open circles indicate stress granules in HeLa_CTG_960__GFPMBNL1 cells (n=25). All data displayed as mean+/- SD.

### MBNL1 dynamics within stress granules show subtle alterations in HeLa cells containing CUGexp foci

To investigate the effect of CUGexp foci on the dynamics of MBNL1 within stress granules, FRAP analysis was carried out on HeLa_CTG_12__GFPMBNL1 and HeLa_CTG_960__ GFPMBNL1 cells induced with Dox for 24hrs prior to treatment with sodium arsenite for 45mins (Fig.5C and D). This analysis revealed a smaller mobile fraction for GFPMBNL1 (28.7%+/-9.5 for HeLa_CTG_12__GFPMBNL1 and 34.8%+/-16.2 for HeLa_CTG_960__ GFPMBNL1) in stress granules to that seen in HLECs (Fig.2), and a slightly shorter halftime of recovery (1.8s+/-1.1 for HeLa_CTG_12__GFPMBNL1 and 2.2s+/11.7 for HeLa_CTG_960__ GFPMBNL1), suggesting GFPMBNL1 is less able to exchange within stress granules in HeLa cells compared to lens epithelial cells. No statistically significant differences in GFPMBNL1 FRAP data between the two HeLa cell lines, that could be attributed to the presence of CUGexp expansion, were detected using a Mann-Whitney test (Fig.5C). Visualisation of individual data sets, however, suggests increased variability in the data obtained for line HeLa_CTG_960__ GFPMBNL1 compared to the control cell line, with the cell line containing CUGexp repeats showing an apparent split of t1/2 measurements into two groups. This idea was supported by an increased variance in the t1/2 values for the CTG960 line in comparison to the CTG12 line (76.5% cf 59.6%). Furthermore, a D’Agostino-Pearson omnibus normality test also confirmed that the half-times of recovery of GFPMBNL1 in Stress granules in line HeLa_CTG_960__ GFPMBNL1 do not show a normal distribution P=0.0084, while those for the control line are normally distributed.

### Stress Granule Dynamics are altered by reduction in expression of MBNL1 or CUGBP1

Having established that MBNL1 and CUGBP1 co-localise closely in stress granules and that there may be subtle changes in stress granule structure and dynamics associated with the presence of CUGexp nuclear foci, we next sought to investigate the relationship between these two DM1-associated proteins in stress granules. Stable cell lines were established containing doxycycline-inducible lentiviral shRNAs against MBNL1 (HeLa_pLKO_MBNL1) or CUGBP1 (HeLa_pLKO_CUGBP1). Robust reduction of MBNL1 (Fig. 6A) or CUGBP1 (Fig. 6B) following doxycycline induction was validated using both immunocytochemistry and immunoblotting. FRAP analysis of transiently-expressed GFPCUGBP1 in line HeLa_pLKO_MBNL1 and GFPMBNL1 in line HeLa_pLKO_CUGBP1 was then used to compare the dynamics of the two proteins in stress granules and to assess the effects of the reciprocal knock-downs.

**Figure 6.**
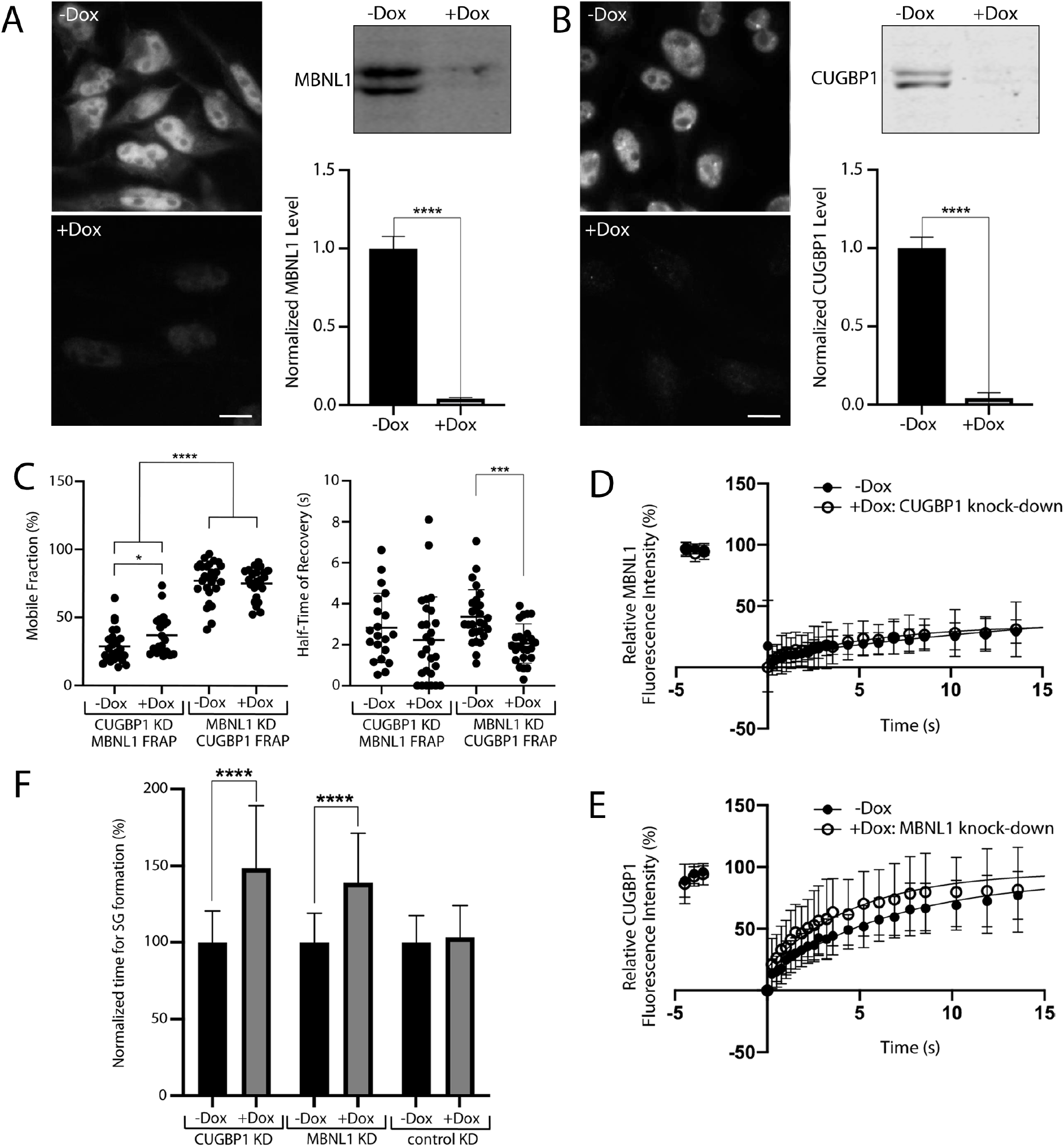
Stress Granule Dynamics are altered by reduction in expression of MBNL1 or CUGBP1. A) Doxycycline-inducible reduction of MBNL1 validated using antibodies against endogenous MBNL1 for immunocytochemistry (left) and immunoblotting (right). Bar =10μm, ****P<0.0001 (data from 3 independent experiments). B) Doxycycline-inducible reduction of CUGBP1 validated using antibodies against endogenous CUGBP1 for immunocytochemistry (left) and immunoblotting (right). Bar =10μm, ****P<0.0001 (data from 3 independent experiments). C) Graphs showing the Mobile Fraction (left) and Half-Time of fluorescence Recovery (right) of GFPMBNL1 and GFPCUGBP1 in stress granules with (+Dox) and without (-Dox) knockdown (KD) of either CUGBP1 or MBNL1. **** P<0.0001, * P<0.05, ***P<0.001 D) Curve fit for FRAP of GFPMBNL1 in Stress granules in cells with CUGBP1 knockdown (+Dox, closed circles, n=21) compared to controls (-Dox, open circles, n=27). E) Curve fit for FRAP of GFPCUGBP1 in stress granules in cells with MBNL1 knockdown (+Dox, closed circles, n=24) compared to controls (-Dox, open circles, n=27).F) Bar Chart showing the time taken for stress granule formation in cell lines with Dox-inducible CUGBP1 and MBNL1 knock-down compared to a negative control, normalised to the mean values for each cell line without knock-down (-Dox). ****P<0.0001 (for CUGBP1 KD -DOX: n=96; +DOX: n=107; for MBNL1 KD -DOX: n=56; +DOX: n=60; for control (luciferase) KD -DOX: n=20; +DOX: n=14). All data displayed as mean+/-SD.

In un-manipulated cells, CUGBP1 had a significantly higher mobile fraction than MBNL1 (77.2% +/-14.5 for CUGBP1 cf 28.8% +/-12.3 for MBNL1), but a comparable half-time of recovery (compare data sets -Dox in Fig.6C). Reduction of MBNL1 expression resulted in a decreased half-time of recovery for GFPCUGBP1, without altering the mobile fraction, indicating that the same amount of CUGBP1 was free to exchange within the stress granules, but that its rate of movement was faster (Fig.6CandE). Reduction in CUGBP1 expression resulted in a slight increase in the mobile fraction for GFPMBNL1, without altering the halftime of recovery, indicating that a larger amount of GFPMBNL1 was free to exchange within the stress granules but with the same rate of movement (Fig.6CandD). This suggests a complex relationship between MBNL1 and CUGBP1 in stress granules, with each protein having some effect on the dynamics of the other.

Time-lapse microscopy was then used to assess the kinetics of stress granule formation in response to sodium arsenite treatment in in HeLa cells with reduced expression of MBNL1 or CUGBP1 (Fig.6F). For both proteins, reduction of expression using doxycycline treatment resulted in a delay in stress granule formation. This delay was not seen in controls using expression of shRNAs targeting luciferase. For MBNL1 knock-down, the delay in stress granule formation was also markedly less (38.9% +/-32.4%) than that seen following CUGexp induction (141.6%, +/-81.1) (Fig.5B), despite the high efficiency of MBNL1 reduction.

### P-Bodies and Stress granules both behave as liquid-liquid-phase-separated (LLPS) structures in Lens Epithelial cells

In mammalian cells, both stress granules and P-bodies are thought to be examples of liquidliquid-phase-separated (LLPS) structures, formed as a result of weak, promiscuous interactions between low-complexity prion-like domains (PrLDs) found in their key protein components (Jain et al., 2016; Kroschwald et al., 2015; Protter and Parker, 2016). The rapid exchange of MBNL1 and CUGBP1 detected by FRAP (Figs 2,5 and 6) within stress granules and P-bodies and the ability of stress granules to fuse and separate (Fig.5 and supplementary movies 1-3) seen in our two models of DM1 are consistent with this hypothesis. Furthermore, analysis of the amino acid sequences of CUGBP1 and MBNL1 using PLAAC (Prion-Like Amino Acid Composition), software (Lancaster et al., 2014) designed to predict prion-like regions, revealed likely prion-like domains in both proteins, with MBNL1 having a longer region in its sequence than CUGBP1 (Fig.7A).

**Figure 7.**
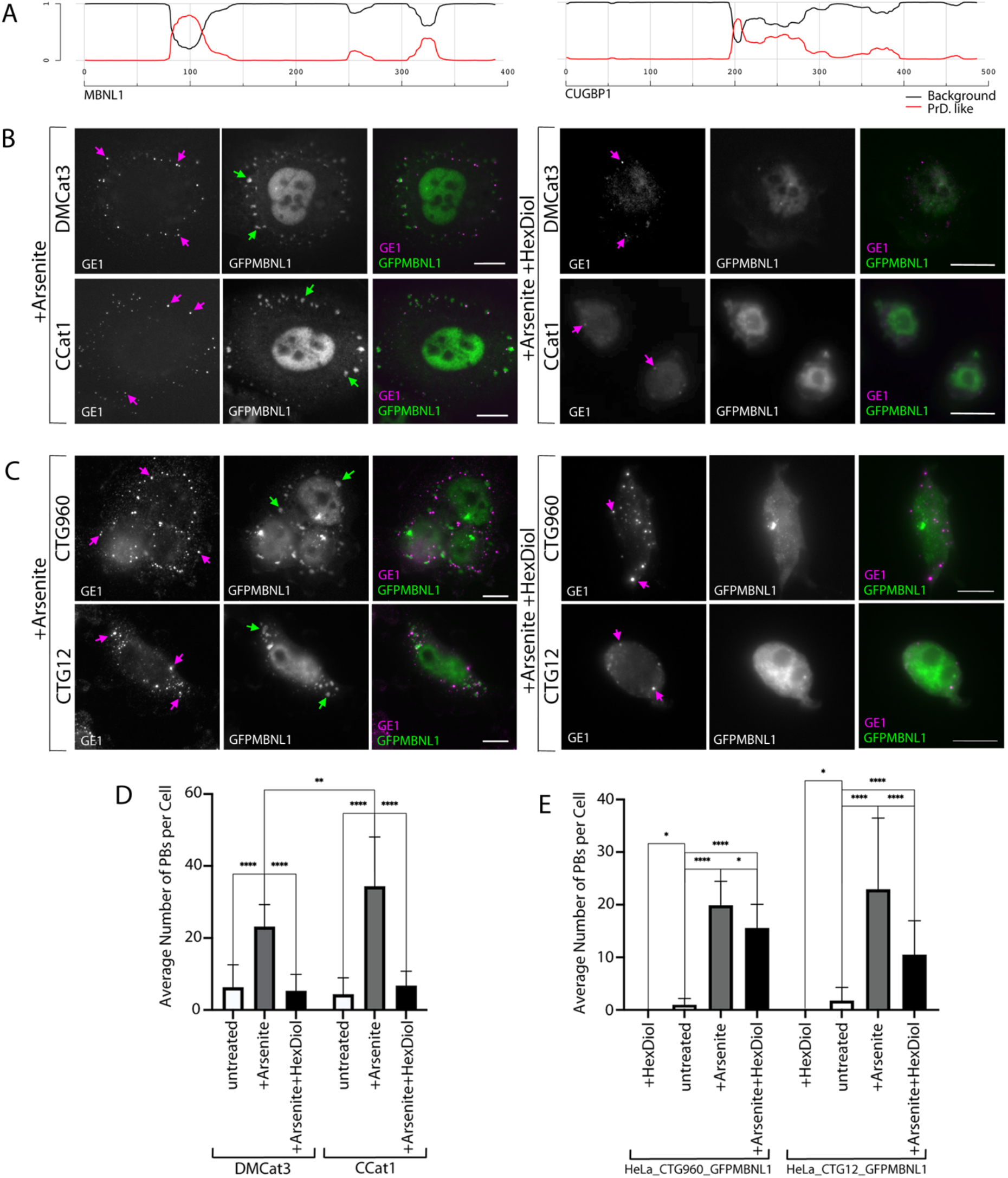
P Bodies and stress granules both show characteristics of liquid-liquid-phase-separated (LLPS) structures, with PB stability increased by the induction of CTGexp repeats. A) Graphical representation of the PLD prediction for MBNL1 (left) and CUGBP1 (right). The output was generated using the Prion-like Amino Acid Composition (PLAAC) program, found at http://plaac.wi.mit.edu/ (Lancaster et al., 2014). The parameters used were “core length” of 30 and the “homo sapiens” background frequency. Prion-like domains are depicted where the probability is >0.5 on the Y axis (red line, PLD domain probability score; black line, “background” probability score). B) DM1 (top row) and control (bottom row) human lens epithelial cells treated with sodium arsenite (left) and sodium arsenite followed by 1,6-hexanediol (right). Stress granules (green arrows) containing GFPMBNL1 (green on overlay) are clearly seen in arsenite treated cells and are absent following treatment with 1,6-hexanediol. P-bodies (magenta arrows) detected with antibodies to endogenous GE1 (magenta on overlays) are numerous in arsenite treated cells and are less numerous following treatment with 1,6-hexanediol. C) Cells from line HeLa_CTG_960__MBNL1-GFP treated with sodium arsenite (left) and sodium arsenite followed by 1,6-hexanediol (right). Stress granules (green arrows) containing GFPMBNL1 (green on overlay) are clearly seen in arsenite treated cells and are absent following treatment with 1,6-hexanediol. P-bodies (magenta arrows) detected with antibodies to endogenous GE1 (magenta on overlays) are numerous in arsenite treated cells and are less numerous following treatment with 1,6-hexanediol. Bar=10μm D) P-bodies per cell in DM1 (DMCat3) and control (CCat1) HLECs treated with 0.5mM sodium arsenite and treated with 0.5mM sodium arsenite followed by 5% 1,6-Hexanediol compared to untreated cells. E) P-bodies per cell in HeLa cells with (HeLa_CTG_960__MBNL1-GFP) and without (HeLa_CTG_12__MBNL1-GFP) CTGexp foci treated with 0.5mM sodium arsenite and treated with 0.5mM sodium arsenite followed by 5% 1,6-Hexanediol compared to untreated cells. ****P<0.0001, ***P<0.001, **P<0.01,*P<0.05. For DMCat3 n=53; for CCat1 n=53; for HeLa_CTG_960__GFPMBNL1 n=108; for HeLa_CTG_12__GFPMBNL1 n=69 All data displayed as mean +/-SD.

Having identified subtle changes in the structure, contents and dynamics of stress granules and P-bodies associated with the presence of CUGexp foci, we next sought to investigate their behaviour in response to the solvent 1,6-Hexanediol, previously shown to disrupt LLPS structures in yeast and mammalian cells (Kroschwald et al., 2015). To determine whether stress granules and P-bodies behave as liquid-liquid phase-separated (LLPS) structures in human lens epithelial cells (HLECs), cells were pre-treated with sodium arsenite, then treated with 5% 1,6-hexanediol for 20 minutes. Stress granules, detected using antibodies against endogenous MBNL1, dispersed on 1,6-Hexanediol treatment of DM1 and control HLECs (Fig.7B), indicating that stress granules in these cells demonstrate the liquid-like property of dispersal following 1,6-hexanediol treatment in addition to the rapid exchange of their protein components and their ability to fuse and split. This is consistent with analysis of the biophysical properties of stress granules in other mammalian cell lines. For P-bodies, detected with antibodies against endogenous GE1, the initial treatment with sodium arsenite significantly increased their numbers in both DM1 and control cells (Fig.7B and D). The numbers of P-bodies seen following stress induction were significantly higher in DM1 cells compared to controls, despite being comparable in unstressed cells (Fig.7D). Subsequent treatment with 1,6-Hexanediol resulted in a marked reduction in the number of P-bodies per cell back to a level comparable to that seen in unstressed cells in both DM1 and control cells (Fig.7B and D). This suggests that the majority of P-bodies in HLECs subject to arsenite stress are formed by LLPS, but that a few P-bodies may represent more stable structures showing insensitivity to 1,6-Hexanediol treatment.

### P-bodies induced by sodium arsenite in HeLa cells containing CTGexp repeats are resistant to treatment with 1,6-hexanediol

Further investigation of the characteristics of P-bodies in stressed cells was made using the inducible HeLa cell model, to allow a more direct assessment of any effect of the presence of CUGexp foci in a genetically homogeneous background. Exposure of cells from lines HeLa_CTG_12__GFPMBNL1 and HeLa_CTG_960__GFPMBNL1 to 1,6-Hexanediol without pretreatment with sodium arsenite resulted in the dispersal of all P-bodies in both lines compared to untreated cells, demonstrating liquid-like properties for P-bodies in unstressed cells (Fig.7E). Treatment with sodium arsenite resulted in an increased number of P-bodies, similar to that seen in HLECs. Subsequent treatment with 1,6-Hexanediol significantly reduced the number of P-bodies in both cell lines, but in contrast to the results from HLECs, the number of P-bodies per cell did not return to pre-stressed levels (Fig.7CandE). Notably, the average number of P-bodies per cell remained much higher in cells containing CUGexp foci than in those without. Taken together, these data suggest that the P-bodies present in arsenite stressed cells represent a diverse population with differing degrees of sensitivity to 1,6-hexanediol. Furthermore, the increase in 1,6-Hexanediol resistant P-bodies in HeLa_CTG_960__GFPMBNL1 suggests that the presence of the CUGexp foci pre-dispose the cells to the formation of more stable P-bodies when subjected to stress.

## Discussion

### The DM1-associated proteins MBNL1 and CUGBP1 are both components of P-bodies and of stress granules

In DM1, the activities of the two multi-functional RNA-binding proteins, MBNL1 and CUGBP1 are known to be altered. MBNL1 is recruited to aberrant nuclear CUGexp foci, which are the hallmark of DM1, while the activity of CUGBP1 is upregulated. There is evidence that this leads to changed patterns of alternative splicing in DM1, with MBNL1 and CUGBP1 acting antagonistically as splicing regulators (de Haro et al., 2006; Konieczny et al., 2014; Pettersson et al., 2015; Wang et al., 2015). The full extent of the effects of DM1 on RNA production and turn-over within the cell, however, are not understood.

Following our observation that MBNL1 and CUGBP1 co-localise in small punctate cytoplasmic structures, despite showing no obvious co-localisation within the nucleus, we investigated the localisation and dynamic behaviour of MBNL1 and CUGBP1 in the cytoplasm in two independent cell culture models of DM1. In Human lens epithelial cells (HLECs) from DM1 patients and controls (Fig.1) MBNL1 and CUGBP1 co-localise in P-bodies in cells grown under normal conditions and in and stress granules following sodium arsenite treatment to induce cellular stress. P-bodies and stress granules are closely related structures involved in cytoplasmic mRNA regulation, and there is some evidence that components of P-bodies re-locate to stress granules in stressed cells (Anderson et al., 2015; Protter and Parker, 2016; Shah et al., 2016). In our novel inducible HeLa cell model of DM1 (Figs.3 and 4), the situation is slightly more complex. CUGBP1 and MBNL1 co-localise in numerous cytoplasmic stress granules following arsenite treatment but in un-stressed cells, whereas CUGBP1 accumulates in P-bodies regardless of the presence of CUGexp foci, MBNL1 appears to be absent from P-bodies in cells with CUGexp foci in their nuclei. This observation suggests that the role of MBNL1 in P-bodies may be disrupted by the presence of CUGexp RNA.

### No underlying basal stress response is detected in human lens epithelial cells from DM1 patients or in an inducible HeLa cell model of DM1

In contrast to previous reports in DM1 myoblasts compared to controls under normal growth conditions (Huichalaf et al., 2010; Ravel-Chapuis et al., 2016) suggesting that DM1 leads to a higher basal level of stress, we did not see evidence of stress granules in either of our cell models grown under standard culture conditions, but only when subjected to stress. The reason for this is unclear, but the difference is unlikely to be a simple correlation with the size of the CUGexp foci which, although small in HLECs from DM1 patients (Coleman et al., 2014) are large in our inducible HeLa cell model. The difference may reflect the use of different cell types and highlights the complexity of the cellular pathology of DM1, which is a multi-systemic condition with highly variable symptoms. The lens of the eye is highly vulnerable to cataract development in DM1 patients (Smith and Gutmann, 2016), but an underlying basal stress response in lens epithelial cells, which act as stem cells for lens development throughout life appears unlikely to be the cause. It is possible, though, that an underlying increased basal stress response may be present in cells of the lens at later stages of differentiation, leading to cataract formation.

### Stress granule turn-over is altered in cell culture models of DM1

Analysis of HLECs treated with sodium arsenite to induce oxidative stress revealed no increased rate of formation of stress granules in cell lines derived from DM1 patients compared to age-matched controls (Fig.2). In our novel HeLa cell model, which contains larger CUGexp foci than are seen in DM1 patient-derived HLECs, a pronounced delay in stress granule formation was seen (Fig.5). Impaired stress granule formation, in the absence of a detectable underlying stress response, has been reported previously in fibroblasts from DM1 patients induced to adopt a myoblast phenotype by exogenous expression of MyoD (Ravel-Chapuis et al., 2016) but not in the unmanipulated fibroblasts. This was interpreted as a cell type difference as myoblasts, but not fibroblasts, are affected in DM1. However, the myoblast cells also had large nuclear foci compared to the parental fibroblasts. Our new results are in broad agreement with these observations, suggesting that there may be a complex interplay between cell type and size of CUGexp foci affecting their response to stress.

The cellular stress caused by sodium arsenite is reversible on its removal. In contrast to the delay seen in formation of stress granules only in the HeLa cell model, both of our cell models showed an increased rate of stress granule dispersal in cells containing the CUG expansion following the removal of stress. In the inducible HeLa cell model, this effect was more pronounced in cells with CUGexp foci induced for longer periods of time (72 hrs cf 24 hrs), suggesting that the changes are directly linked to the formation and persistence of the foci. Stress granules are transient structures, with the mRNAs sequestered within them destined to return to active translation. The dispersal of stress granules on the removal of stress is a tightly regulated process involving a form of autophagy (Hardy et al., 2017; Mateju et al., 2017; Protter and Parker, 2016; Turakhiya et al., 2018). The rapid loss of stress granules from cells containing CUGexp foci may indicate alterations or impairment in the mechanisms required for regulated stress granule dispersal associated with the presence of the CUG expansion. This would require considerable additional analyses to investigate, but may be of importance for understanding the cellular pathology of DM1 in the context of cataract development. Defects in autophagy are associated with cataract development, though the mechanisms and types of autophagy involved are, as yet, unclear (Costello et al., 2013; McWilliams et al., 2019; Morishita et al., 2013; Morishita and Mizushima, 2016).

### Alterations to stress granule turn-over cannot be attributed solely to the sequestration of MBNL1 in CUGexp foci

The sequestration of MBNL1 in CUGexp foci has long been cited as a major cause of cellular defects in DM1. However, we have previously reported that the amount of total cellular MBNL1 present in nuclear foci in HLECs from DM1 patients is extremely small (0.2% (Coleman et al., 2014). Even in our HeLa cell model, which appears to contain large and very bright nuclear foci when viewed using GFPMBNL1, the amount of cellular GFPMBNL1 contained in the foci is only 1.7% (+/-1.4) (Fig. S3). Delayed formation of stress granules was also seen in HeLa cells without CUGexp foci but with shRNA-mediated MBNL1 depletion (Fig.6). While this does implicate loss of MBNL1 as a direct cause of disruption to stress granules formation, the delay seen following MBNL1 reduction by shRNA was much less pronounced than that resulting from CUGexp foci induction, despite the almost complete loss of MBNL1 seen in shRNA-treated cells. Although in DM1 it is likely that MBNL1 is also bound to CUG repeats in DMPK mRNA outside the nuclear foci, the small size of the effect on stress granule turnover of almost complete MBNL1 depletion compared to that seen in cells containing CUGexp foci strongly suggests that sequestration of MBNL1 is not the primary cause of the alterations to stress granule turnover seen.

### MBNL1 and CUGBP1 are likely core components of stress granules and may be recruited from P-bodies

MBNL1 and CUGBP1 have different dynamics within stress granules, with GFP-CUGBP1 showing both a higher mobile fraction and faster recovery in FRAP experiments than MBNL1-GFP (Fig.6). Previous studies of various stress granule proteins in different cell types have reported highly divergent dynamic behaviours (Bley et al, 2015), with proteins essential for stress granule formation exhibiting faster kinetics within stress granules (T1/2 of 2 to 3 seconds) and higher mobile fractions than those dispensable for their formation. MBNL1 exhibits similar behaviour to that published for TIA-1, a well established marker for stress granules whose depletion results in impaired stress granule formation (Bley et al., 2015; Gilks et al., 2004). Together with the delayed formation of stress granules in cells with disrupted MBNL1 expression, this suggests that MBNL1 is important for stress granule integrity. CUGBP1, while exhibiting rapid exchange within stress granules similar to that of MBNL1, has a small mobile fraction previously reported to be characteristic of non-essential stress granule components. Reduction of CUGBP1 expression, however, also causes a delay in stress granule formation, which would not be expected for a non-essential component, emphasising further the complexity of these structures.

It has been suggested that stress granules in mammalian cells are composed of a mixture of liquid and solid phases (Protter and Parker, 2016) with a solid core structure surrounded by a more dynamic shell (Jain et al., 2016; Markmiller et al., 2018; Protter and Parker, 2016). High resolution airyscan microscopy reveals a close co-localisation between MBNL1 and CUGBP1 in stress granules, with both proteins showing punctate accumulations similar to those reported as stress granule cores containing GFP-G3BP detected by STORM microscopy (Jain et al., 2016). CUGBP1 was also previously identified in the proteome of stress granule core structures (Jain et al., 2016; Kuechler et al., 2020; Youn et al., 2018). This is slightly at odds with the small mobile fraction we see for CUGBP1 in FRAP analyses, which would suggest it is a non-essential component, but overall strengthens the evidence for the two DM1-associated proteins MBNL1 and CUGBP1 as core stress granule proteins. This represents a slightly unusual situation, as proteins required for the formation of a particular RNP structure are usually confined to that structure (reviewed in (Corbet and Parker, 2019)), while MBNL1 and CUGBP1 are also found in P-bodies as well as in specific regions of the nucleus (Coleman et al., 2014; Fujimura et al., 2008). However, P-bodies and stress granules show significant overlap in their components (reviewed in (Youn et al., 2019)) and it has been known for some time that stress granules and P-bodies are closely linked structures that can dock and fuse with each other (reviewed in (Stoecklin and Kedersha, 2013).

In our inducible HeLa cell model of DM1, the presence of CUGexp RNA resulted in a slight decrease in Pearson’s co-efficient of co-localisation between GFPMBNL1 and CUGBP1 within stress granules, suggesting a slight alteration in the fine structure of stress granules. In the same cells, both CUGBP1 and GFPMBNL1 were detected in P-bodies in un-stressed cells without CUGexp repeats, but MBNL1 was absent from P-bodies in cells containing CUGexp repeats. Following induction of stress using sodium arsenite, GFPMBNL1 was seen in a subset of P-bodies in both cell lines, but the amount of GFPMBNL1 in P-bodies was significantly reduced in cells with CUGexp foci. It is conceivable that this loss of MBNL1, as a core stress granule component, from P-bodies contributes to the delayed formation of stress granules seen in these cells following treatment with sodium arsenite.

### Alterations to liquid-liquid phase separation (LLPS) may be associated with DM1

Prion-like domains within low-complexity sequences promote the formation of LLPS structures, including mammalian stress granules (Ditlev et al., 2018; Markmiller et al., 2018; Protter & Parker, 2016), as it is these domains that form the weak promiscuous interactions necessary for phase separation to occur. Analysis of the sequences of MBNL1 and CUGBP1 using the PLAAC (Prion-Like Amino Acid Composition) web application (Lancaster et al., 2014) to predict prion-like amino acid compositions, reveals a small predicted prion-like domain in both MBNL1 and CUGBP1, which may account for their propensity to accumulate Stress granules. The domain in MBNL1 is longer (28 amino acids) than that in CUGBP1 (10 amino acids).

The induction of CUGexp RNA in our HeLa cell model of DM1 resulted in a subtle difference in the dynamics of MBNL1 within stress granules. While no statistically significant change was seen in the overall dynamics of MBNL1 within stress granules, the distribution of values for the half-time of recovery showed a subset of stress granules with a much slower rate of return of MBNL1 following CUGexp foci induction. The explanation for this is unclear but this observation could be explained by an increased tendency for the formation of more solid stress granules in cells containing nuclear foci. However, in cells with MBNL1 reduced to very low levels using shRNA (Fig.6), we observed an increase in the rate of exchange of CUGBP1 within stress granules. If the rate of protein exchange can be taken as a direct indication of the fluidity of a structure this would suggest that MBNL1 reduction, predicted to result from its sequestration in CUGexp foci, decreased the solidity of stress granules. This apparent contradiction, taken together with the differences in rates of exchange of MBNL1 and CUGBP1 in stress granules despite their close co-localisation, suggest that the dynamics of LLPS structures are extremely complex (reviewed in (A and Weber, 2019) and that FRAP is not a perfect method to examine the biophysical nature of all cellular structures.

1,6-Hexanediol has been extensively used to disrupt structures predicted to form by LLPS, *in vitro* and *in vivo* (Kroschwald et al., 2015; Molliex et al., 2015; Patel et al., 2007; Ribbeck and Gorlich, 2002; Updike et al., 2011). Treatment of human lens epithelial cells and inducible HeLa cell models of DM1 with 1,6-hexanediol resulted in complete loss of stress granules, as would be expected for a phase-separated structure. The effect on P-bodies, however, was more complex. In un-stressed HeLa cells, no P-bodies were visible after treatment with 1,6-hexanediol. However, in HeLa cells treated with arsenite before 1,6-hexanediol, large numbers of additional P-bodies were formed during the stress response and a proportion of these remained following 1,6-hexanediol treatment, suggesting that they had lost their liquid-like properties. Of particular interest for understanding DM1, the number of 1,6-hexanediol-resistant P-bodies was significantly higher in HeLa cells containing CUGexp nuclear foci. This suggests that the presence of the DM1-associated CUG expanded RNA influences the biophysical characteristics of P-bodies following cellular stress.

### Implications of altered SG and PB dynamics for DM1 pathology

Defects in the normal physiology of cytoplasmic LLPS, particularly stress granules, have been widely implicated in human diseases. The spontaneous conversion of prion-like domains into an aggregated state, and subsequent transition from liquid to solid to form solid RNP aggregates has been proposed as a pathological mechanism in conditions including ALS and FTD (Li et al., 2013; Ramaswami et al., 2013). The presence of aberrant or persistent stress granules is also associated with the cellular pathology of age-related diseases, neurodegeneration and some cancers (Mateju et al., 2017; Protter and Parker, 2016; Turakhiya et al., 2018). Furthermore, increased basal stress levels and defects in cellular stress responses have been suggested to occur in DM1 (Huichalaf et al., 2010; Ihara et al., 1995; Kumar et al., 2014; Toscano et al., 2005; Usuki and Ishiura, 1998; Usuki et al., 2000). Here we report delayed stress granule formation and premature stress granule loss in cell culture models of DM1, together with altered biophysical characteristics of the closely related P-bodies. This strongly suggests that the presence of CUGexp RNA, perhaps partly as a consequence of the sequestration of MBNL1 by the CUG expansion within foci in the nucleus of the cell or elsewhere, results in defects in cellular responses to stress. The assembly and disassembly of stress granules are complex, regulated processes (Wheeler et al., 2016). The failure to disassemble stress granules in a controlled manner could impair the ability of cells to respond effectively to repeated stress events, and furthermore may suggest a defect in autophagy associated with DM1. Defects in autophagy have recently been implicated in several degenerative conditions (Ranganathan et al., 2020) and autophagy is known to contribute to the generation of optically transparent lens fiber cells during the differentiation of the lens epithelial stem cell population (Costello et al., 2013; Morishita et al., 2013; Morishita and Mizushima, 2016).

In summary, our data show alterations to the structure and regulation of stress granules and P-bodies: two key LLPS structures involved in cytoplasmic mRNA metabolism and cellular stress responses in cell culture models of DM1. These changes are associated with the presence of CUGexp RNA and may be linked to deficiency of MBNL1 resulting from its accumulation in the nuclear CUGexp foci characteristic of DM1. However, with only small amounts of MBNL1 co-localising with nuclear CUGexp foci, and the failure of almost complete MBNL1 knock-down to recapitulate the magnitude of the effects seen in cells expressing CUGexp RNA, MBNL1 sequestration in CUGexp foci is unlikely to provide a full explanation. Our observations point to a complex interplay between nuclear and cytoplasmic structures involved in mRNA metabolism, and suggests that further investigation of the contribution of both altered LLPS and autophagy to cellular defects in DM1 is warranted.

## Acknowledgements

This work was supported by Muscular Dystrophy UK (project grant number 16GRO-PG36-0065 to JES), BBSRC (EASTBIO DTP studentship award to SG) and the Wellcome Trust (ISSF award 105621/Z/14/Z to JES and PG). Plasmids pBI-Tet-CTG12 and pBI-Tet-CTG960 were a gift from Prof. T Cooper, Baylor College of Medicine, Houston. Plasmid Tet-pLKO-puro_shLUC was a gift from Dr Christina Paulus, University of St Andrews, UK, who also assisted with FACS sorting during generation of the inducible HeLa cells lines. Antibody Mbla was a gift from Prof. Glenn Morris, Wolfson Centre for Inherited Neuromuscular Disease (CIND), The Robert Jones and Agnes Hunt Orthopaedic and District Hospital, Oswestry, UK.

