## Supplemental figures and legends for "Phase-separated stress granules and processing bodies are compromised in Myotonic Dystrophy Type 1"

**Supplementary Data**

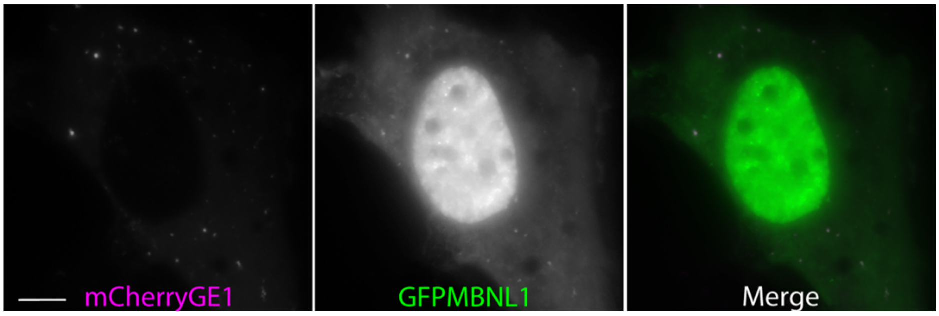


***Figure S1 (related to figure 2): Verification of PB identity in living cells prior to FRAP analysis.*** *Human lens epithelial cells were transiently transfected with plasmids to express GFPMBNL1 (centre and green on merge)and mCherryGE1 (left and magenta on merge), a marker for P-bodies. Cytoplasmic foci conclusively identified as P-bodies using mCherryGE1 were targeted for bleaching in FRAP experiments. Bar = 5μm.*

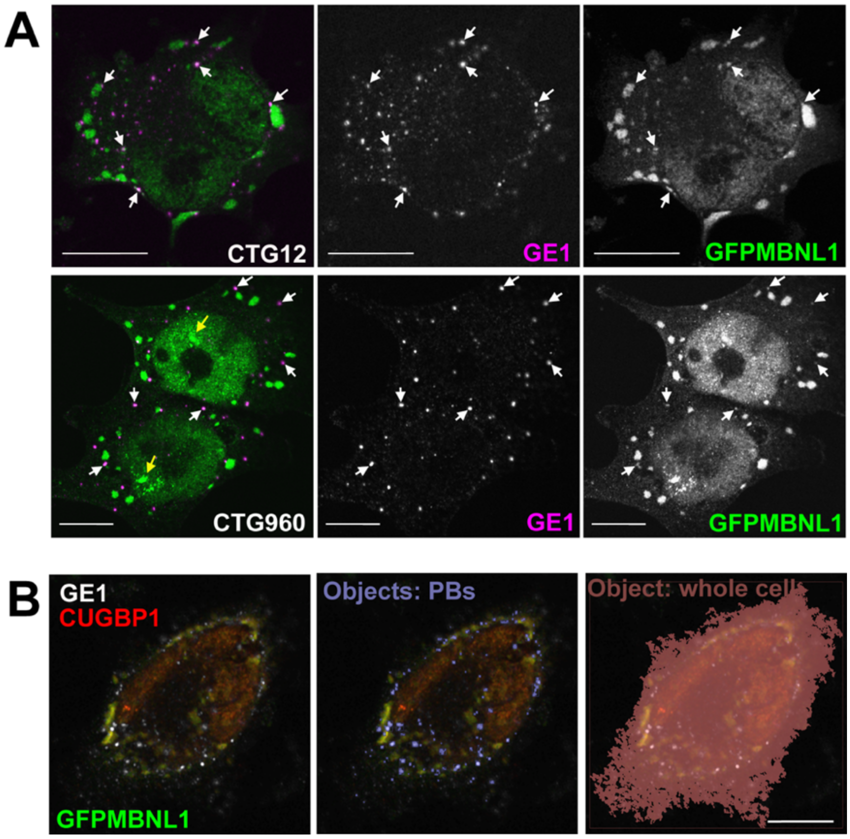


***Figure S2 (related to figure 4): Quantitation of GFPMBNL1 in P-bodies in HeLa cell model of DM1****. A) Cells from line CTG12 (top) and CTG960 (bottom) treated with sodium arsenite contain large numbers of P-bodies detected with antibodies against GE1 (centre and magenta on merge) in addition to Stress granules. Some, but not all, of these P-bodies contain detectable amounts of GFPMBNL1 (right and green on merge), examples marked by white arrows. Yellow arrows denote CUGexp nuclear foci containing GFPMBNL1. B) Example of object identification used to calculate the % of total cellular GFPMBNL1 per PB. Signals from the original images (left) were used to identify P-bodies using GE1 (centre, purple objects) and the entire cell volume using CUGBP1 (right, brown object). Bar=10μm.*

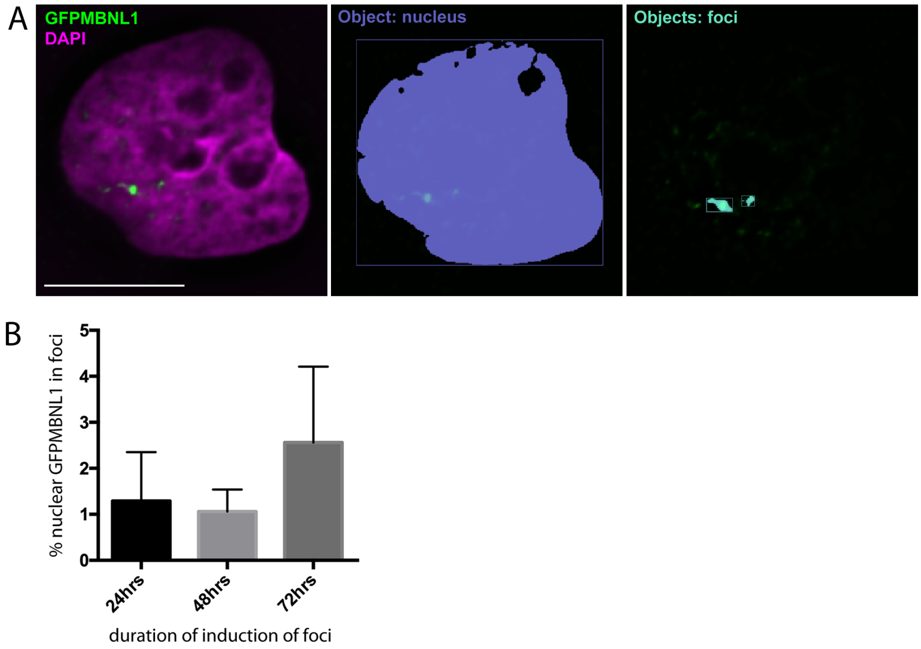


***Figure S3 (related to discussion): Quantitation of GFPMBNL1 in nuclear CUGexp foci in HeLa cell model of DM1.*** *Example of object identification used to calculate the % of total nuclear GFPMBNL1 in nuclear CUGexp foci. Signals from the original images (left) were used to identify foci using GFPMBNL1 (right, cyan objects) and the nuclear volume using DAPI (centre, purple object). Bar=10µm.*

***Movie S1: SG formation (CTG12)***
*Maximum intensity z-projection of time-lapse movies showing the formation of cytoplasmic stress granules in cells from line CTG12 treated with sodium arsenite (related to figure 5).*

***Movie S2: SG formation (CTG960)***
*Maximum intensity z-projection of time-lapse movies showing the formation of cytoplasmic stress granules in cells from line CTG960 treated with sodium arsenite (related to figure 5).*

***Movie S3: SG dispersal***
*Maximum intensity z-projection of time-lapse movies showing the loss of cytoplasmic stress granules from cells from line CTG960 following sodium arsenite removal. The splitting of Stress granules into smaller structures prior to their complete loss can be seen (related to figure 5).*
